# Simultaneous neuron evidence for much higher covariation with saccadic reaction time of superior colliculus than primary visual cortex visual responses

**DOI:** 10.64898/2026.05.19.726219

**Authors:** Yue Yu, Ziad M. Hafed

## Abstract

Visual response strength in the primate superior colliculus (SC) has recently been shown to inversely correlate with trial-by-trial saccadic reaction time in a much stronger way than visual response strength in the primary visual cortex (V1). However, for any given visual stimulus onset, populations of neurons in each brain area are concurrently activated, leaving open the question of how V1 visual response strength can predict trial-by-trial saccadic reaction time when multiple simultaneously recorded neurons are taken into account. Using a classic visually-guided saccade task, here we assessed the quality of predicting trial-by-trial saccadic reaction time from the visual response strengths of 1 to 10 simultaneously recorded neurons in each brain area. For each session, we modeled saccadic reaction time as a weighted linear combination of the visual response strengths of *N* simultaneously recorded neurons. Consistent with the prior work, the visual response strength of a single SC neuron was better than that of a single V1 neuron at predicting reaction time. By adding more simultaneously recorded neurons, the prediction got much better in the SC, but not in V1. Only for 100% contrast dark stimuli (darker in luminance than the surrounding gray background) did V1 show an increase in prediction quality with more simultaneously recorded neurons. This increase, which was still substantially weaker than in the SC, could reflect the ecological relevance of dark contrasts in scenes. These results suggest that despite qualitative similarities between SC and V1 visual responses, SC visual responses are functionally reformatted from their V1 counterparts.

**Significance:** The superior colliculus (SC) is an important sensory-motor structure for controlling eye movements, and it receives a significant portion of its inputs directly from the primary visual cortex (V1). Despite this, SC visual responses are much better correlated with trial-by-trial variability in saccadic eye movement timing than V1 visual responses, and this effect is strongly amplified when considering simultaneously recorded neurons. Thus, SC and V1 visual responses serve fundamentally different functions from a motor perspective.

## Introduction

Saccades are an integral component of active vision, allowing quick alignment of our fovea with regions of interest in our visual environment. These eye movements are known to exhibit substantial levels of variability in their timing (1–7), leading to investigations of the sources underlying such variability. Among these sources is the sensing process itself. Specifically, using different variations of laboratory-designed visually-guided saccade tasks, it was asked whether trial-by-trial variability in the visual response properties (such as latency or strength) of an individual visually-sensitive neuron could be correlated, at least to some extent, with the trial-by-trial variability of saccadic reaction time. In two brain areas that are particularly relevant for eye movement generation, the primary visual cortex (V1) (8) and the superior colliculus (SC) (9, 10), this was indeed the case. However, due to differences in task designs and stimulus properties across the studies, it was difficult to assess which brain area’s visual responses were more closely aligned with behavioral variability than the other’s. This is an important question to resolve, especially because it can help clarify the differential roles of multiple parallel pathways that may connect the retinal visual input to the final oculomotor output. Moreover, resolving this question can aid in understanding the detailed differences that may exist between otherwise similar short-latency visual responses in both the SC and V1.

Recently, we aimed to resolve the above question by measuring SC and V1 visual responses in the same animals and with the very same behavioral tasks and visual stimuli; we found that SC visual responses, and particularly their strengths, were significantly better correlated with saccadic reaction time variability than V1 visual responses (11). However, one limitation of this prior study (11) is that it only considered a single individual neuron in either the SC or V1 at a time. In reality, a single visual stimulus onset would concurrently activate many neurons in each brain area at the same time. Thus, it might be asked whether considering multiple simultaneously recorded V1 neurons might lead to better alignment of V1 visual responses with behavioral variability or not. This is what we investigated here. We related saccadic reaction time variability within a given session to the trial-by-trial visual activity of up to ten simultaneously recorded SC or V1 neurons. Our approach was to use as minimalist a model as possible to predict saccadic reaction time from each brain area’s visual responses. Thus, we expressed reaction time on any given trial as a weighted linear sum of the visual activity of multiple neurons, and we then assessed how well such a simple model could account for across-trial reaction time variability. By using visual response strength as our sensory response property of interest (9–19), we found that adding more simultaneously recorded neurons dramatically improved predicting reaction times in the SC. Remarkably, adding the same number of simultaneously recorded neurons in a V1 model did not improve reaction time prediction by the model, at least in the great majority of conditions. Only when the visual target for saccades was darker than the surrounding background and with 100% Weber contrast did the V1 model clearly improve with increasing numbers of neurons (but still not as strongly as in the case of the SC); this multi-neuron observation deviates from the previous single-neuron analyses (11), which did not suggest that 100% contrast dark stimuli were different than the other stimulus conditions for V1 neurons.

Overall, we found simultaneous neuron evidence for much higher covariation with saccadic reaction time of SC than V1 visual responses. This evidence is important to document for at least three reasons. First, it clarifies the distinction between the SC and V1 when it comes to dictating saccade timing. Specifically, given prior demonstrations of a potential correlation between V1 visual responses and saccadic reaction times (8), it is important to ask how significant such an observation is when compared to the SC, and to also ask whether multi-neuron observations can emerge that could not have been predicted from the previous single-neuron analyses. Second, it demonstrates that the sensing process itself, especially in the SC, is very much a part of the process of saccade generation. Thus, while it is expected that factors like target probability (20, 21), motor preparation (22), as well as SC motor burst generation itself (23, 24) all dictate when a saccade is triggered, all of these processes need to first be jumpstarted by visual responses to stimuli. Finally, comparing the SC and V1 helps support the idea that SC visual responses are not merely inherited from the cortex. This is a fundamental question because it can allow physiological evidence to constrain the space of possible functional interpretations afforded by anatomical facts of connectivity. For example, as we will show below, even though some SC neurons are the biggest recipients of V1 inputs, their relationship to saccade timing is weaker than other SC neurons that have other patterns of anatomical connectivity.

## Materials and methods

### Experimental animals and ethical approvals

The current study involved analysis of the same neural database described recently (11). In that previous investigation, we had used 3 adult, male rhesus macaque monkeys (macaca mulatta). Monkeys A (10-13 years), F (14 years), and M (9 years) were used for the SC recordings; and, monkeys A and F were used for the V1 recordings.

All experiments were approved by ethical committees at the regional governmental offices of the city of Tübingen (Regierungspräsidium Tübingen).

### Laboratory setup

#### The laboratory setup was described in the previous publication (11)

Briefly, experiment control was achieved as follows. We used a modified version of PLDAPS (25) for controlling stimulus presentation and data acquisition; graphics were presented using the Psychophysics Toolbox (26–28). The neurophysiology data acquisition process was achieved with an OmniPlex system from Plexon, and eye tracking was performed using scleral search coils and the magnetic induction technique (29, 30).

Stimuli were displayed on a CRT display of ∼30 by 23 deg dimensions (horizontal and vertical, respectively) and 85 Hz refresh rate. The background was gray (26.11 cd/m^2^), and the different stimuli could have different Weber contrasts relative to this gray value (either brighter or darker than the background). The display was linearized for luminance before the experiments.

Recording for the data that we were analyzing here was performed with linear microelectrode arrays (V-Probes) from Plexon, each having 16, 20, or 24 channels. This enabled recording multiple simultaneous neurons per session.

#### Experimental procedures

The behavioral task for the monkeys was a classic visually-guided saccade task (11). First the monkeys fixated a small (white, 79.9 cd/m^2^) spot of ∼10.8 by ∼10.8 min arc dimensions. After a few hundred milliseconds of fixation, the fixation spot was removed, and an eccentric saccade target appeared simultaneously. The eccentric target was a disc of 0.51 deg radius, and it could have different luminance polarities (darker or brighter than the background). The Weber contrast of the disc from the gray background varied from trial to trial (10%, 20%, 50%, or 100%). The stimulus location was always the same for a given recording session. Thus, the monkeys’ attentional state to the stimulus location was experimentally controlled across all trials.

## Data analysis

For the previous report (11), we detected all relevant eye movement and neural events that were necessary for the present study. Specifically, we detected all saccadic reaction times across trials. Similarly, after spike sorting, we estimated visual response strength for each trial. We did so by counting the number of spikes that occurred in any one trial during a visual response epoch after stimulus onset. The start and duration of the visual response epoch was both area-and contrast-dependent. Specifically, in the SC, the visual response epoch was 60 ms long and starting from 50 ms (for 10% and 20% contrasts) or 40 ms (for 50% and 100% contrasts) after stimulus onset. For V1, the visual response epoch was 85 ms long and starting from 35 ms (for 10% and 20% contrasts) or 30 ms (for 50% and 100% contrasts) after stimulus onset. The contrast-dependence of the visual response epoch reflected the fact that visual responses for lower contrasts tended to be weaker and later than visual responses for higher contrasts (11, 17, 31–35); the area-dependence was because we noticed that visual burst dynamics were different between the two brain areas (11). We do not expect that changing the length of the visual response epoch in a given condition would change the overall results, especially given how transient visual responses can be. Also, we could not extend the integration window too much (forward in time) since there is a subsequent saccade happening within the reaction time period from stimulus onset. In the SC, this subsequent saccade introduces motor bursts (especially in the visual-motor neurons), which we did not intend to include in our analyses.

Our database included sessions with potentially >25 simultaneously recorded neurons. However, since other sessions could have fewer neurons, we decided to limit our analyses to a maximum of 10 simultaneously recorded neurons in any given session. This allowed us to include all sessions in the analyses. Moreover, including more neurons in our model (see below) would require more trial repetitions per condition to avoid overfitting. Thus, 10 neurons was considered to be a suitable number to enable effectively comparing the two brain areas. In total, we had 28 SC sessions and 13 V1 sessions with simultaneously recorded neurons.

From each session (whether in the SC or V1), we collected the saccadic reaction time and visual response strength on each trial of a given stimulus appearance. Then, we found the parameters β*_j_* that provided the best fit to the following equation across all trials with the same visual appearance of the saccade target:

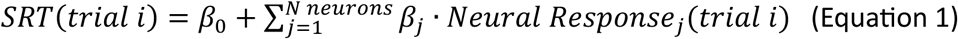

where *SRT(trial i)* is the saccadic reaction time on the *i^th^* trial, *Neural Response_j_(trial i)* is the visual response strength (number of spikes in the visual response epoch) on the *i^th^* trial of the *j^th^* simultaneously recorded neuron in the analysis. To assess the fit quality, we calculated *R^2^_adj_*, after adjusting for different predictor (*p*) and observation (*n*) numbers, using the following formula:

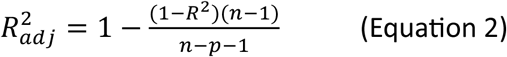

where *R^2^* is the standard coefficient of determination before adjustment.

Within a single session, we plotted *R^2^_adj_* as a function of number of simultaneously recorded neurons. We limited the x-axis (number of simultaneously recorded neurons) to 10, as mentioned above. For sessions with 10 simultaneously recorded neurons, we calculated the *R^2^_adj_* value for each possible combination of *N* neurons (*N* being 1 to 10) that we could obtain within the session. For example, for an *N* value of 2, we calculated *R^2^_adj_* for all possible pairs of two simultaneously recorded neurons within the session. Then, we averaged the *R^2^_adj_* values across all pairs to get the session’s average *R^2^_adj_* value with *N* = 2. We repeated the same procedure for all other values of *N*. For sessions with more than 10 neurons recorded, the number of possible neuron combinations for each value of *N* could increase dramatically (reaching more than 3 million with more than 25 simultaneous neurons within a single session). Thus, we still took combinations of neurons at each value of *N*, but, this time, we only selected the first 1000 permutations of the neurons. This still allowed us to assess how *R^2^_adj_* depended on specific neuron combinations within a given session.

To summarize the population result, we obtained the average *R^2^_adj_* value within each session as mentioned above, and for each *N* value. Then we averaged across all sessions to obtain a population result. We did this for each image luminance polarity and each contrast separately.

Statistically, we performed different comparisons of *R^2^_adj_* at either the individual session level or across the population of sessions. Within each session, we tested whether the mean *R^2^_adj_* values (in a given brain area and for a given contrast level) were statistically different from zero. Specifically, we performed a t-test on the 10 mean *R^2^_adj_* values of each within-session analysis. Then, we performed a regression on the same data to see if the mean *R^2^_adj_* value increased with the number of simultaneously recorded neurons. For the SC, we noticed that the mean *R^2^_adj_* value increased in a nonlinear fashion with neuron number (see Results).

Therefore, the regression was performed after logarithmically transforming the x-coordinate as follows:

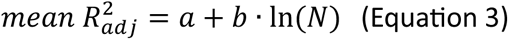

where *a* and *b* are the fit parameters, and *N* is the number of neurons. For V1, the changes with neuron number were linear (if at all), so we did not include the logarithmic transformation of *N* into Equation 3, and simply used a linear regression instead (again, with intercept and slope parameters, *a* and *b*).

Across sessions, for each luminance polarity, we tested whether the *R^2^_adj_* values across contrast levels were statistically different from zero using a t-test. Then, we performed a 3-way ANOVA to check if contrast, number of neurons, and brain area had an effect on the *R^2^_adj_* values. In the end, we also performed an ANOVA on the whole data set, across contrast, luminance polarity, number of simultaneously recorded neurons, and brain area.

We also repeated our analyses of the SC neurons, but after separating these neurons according to whether they were functionally of the visual-motor or visual-only type (36, 37). Visual-motor neurons exhibit saccade-related bursts in addition to their visual responses (as in the example neurons of Fig. 1 in Results); on the other hand, visual-only neurons are the biggest recipients of V1 inputs (19, 38–42). We used the same classification of our neurons that we presented previously (11). Because we essentially had approximately half of the numbers of neurons of each type for this analysis, when compared to the original global database size without functional classification, we restricted our simultaneous neuron analyses to a limit of 5 neurons instead of 10. As we show in Results, this was still sufficient to show the differences between SC visual and visual-motor neurons, as well as the dependencies on increasing numbers of simultaneously recorded neurons.

**Figure 1.**
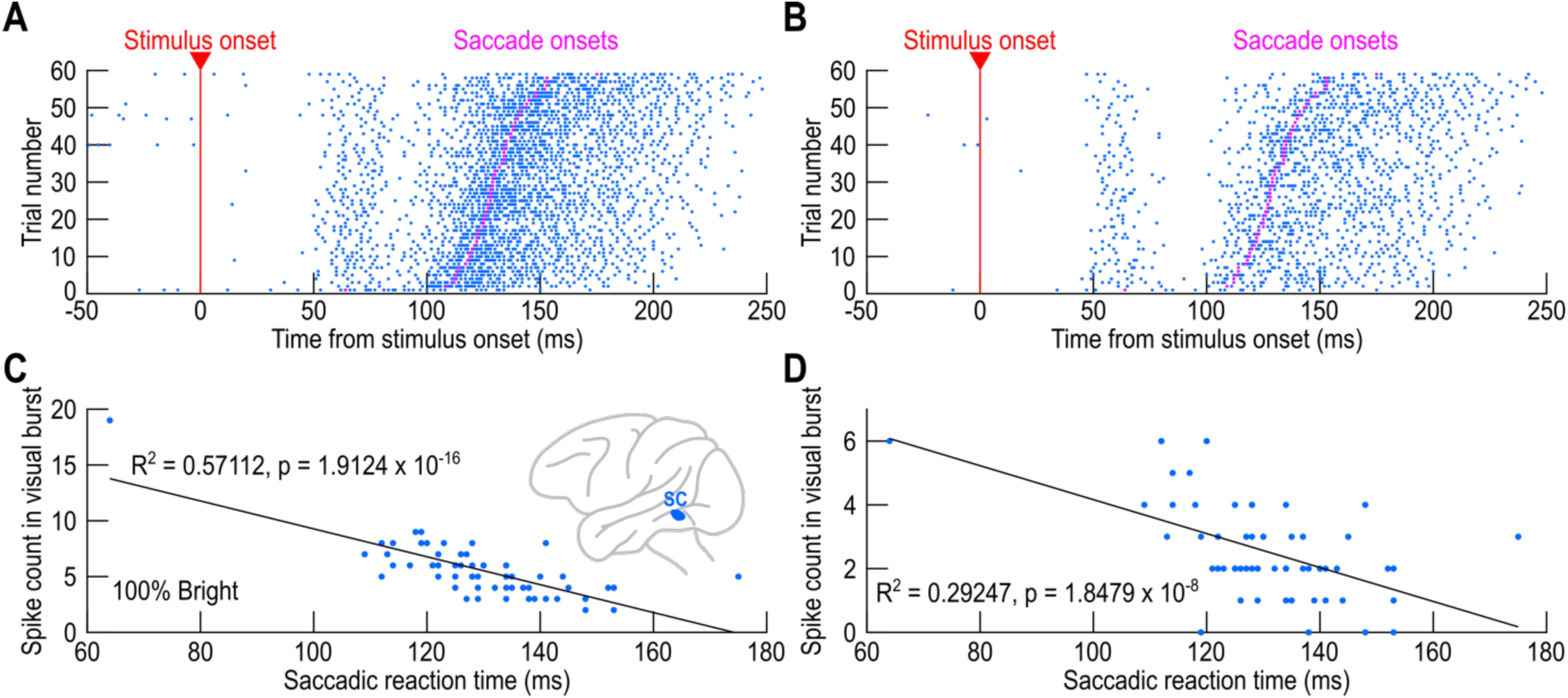
Simultaneously recorded SC neurons exhibited similar negative correlation patterns between trial-by-trial visual response strength and saccadic reaction time. (A, **B)** Spike times of two example simultaneously recorded SC neurons with 100% contrast bright stimuli as the saccade targets. Each row is an individual trial, and each dot is the time of an action potential emitted by the recorded neuron. Both neurons were visual-motor, exhibiting both a visual burst shortly after stimulus onset (at around 50 ms) as well as a saccade-related burst time-locked to saccade onset. The trials were sorted according to saccadic reaction time, and the saccade onsets are indicated by the magenta dots. For both neurons, there were fewer action potentials emitted in the visual burst epoch on trials with long saccadic reaction times than on trials with short saccadic reaction times. (C, D) For each neuron, we plotted the number of action potentials emitted in the visual burst epoch (Materials and methods) as a function of saccadic reaction time. Each dot represents one trial, and the black line is a linear regression fit. The indicated *R^2^_adj_* values are those obtained from Equation 1 with *N* = 1. For both neurons, visual bursts were weaker on slow saccadic reaction time trials. Numbers of trials are evident from the individual rasters in A, B.

Finally, note that we did not analyze visual response onset latencies in the current study. This is because the correlation coefficients to behavior were generally weaker for visual response latency than visual response strength measures (11), and also because we could not always reliably detect a visual response onset latency on every single trial (11). This meant that we could not find enough matching trials with as many simultaneously recorded neurons as with the visual response strength measure, rendering the interpretation of results with visual response latencies harder than visual response strengths.

## Results

### Concurrently active superior colliculus neural populations at stimulus onset are the better predictors of saccade timing

Our goal was to investigate how well we could predict single-trial saccadic reaction times from the visual response strengths of populations of simultaneously recorded SC or V1 neurons, and we did this by employing the simple linear model of Equation 1 (Materials and methods). Using a classic visually-guided saccade task (Materials and methods) (11), we characterized the goodness of fit of Equation 1 within a given session as a function of how many recorded neurons were included into the model.

At the single neuron level (*N* = 1 in Equation 1), our results were consistent with our previous report investigating the same neural database using a slightly different approach (11). For example, Fig. 1 shows an example pair of simultaneously recorded SC neurons from one session. In this case, the 100% Weber contrast bright stimulus was presented within the visual response fields (RF’s) of the two recorded neurons, and we plotted the spike rasters emitted by the two neurons (during the very same trials) after sorting the trials by saccadic reaction time. Both neurons were visual-motor neurons because they emitted both a visual burst aligned to stimulus onset as well as a saccade-related motor burst aligned on saccade onset (Fig. 1A, B). Moreover, both neurons individually showed a negative relationship between saccadic reaction time and visual burst strength. Specifically, Fig. 1C, D shows the number of spikes emitted by each neuron within a visual burst epoch (in the interval 40-100 ms after stimulus onset; Materials and methods) as a function of the subsequent saccadic reaction time observed in the trials. In both cases, weaker visual responses were associated with later saccades (*R^2^_adj_* values from the model of Equation 1 are shown in the figure). In the case of V1, these negative correlations between visual response strengths and saccadic reaction times were much weaker (11), as evidenced by the example pair of simultaneously recorded V1 neurons of Fig. 2 (the visual burst epoch was, in this case, in the interval 30-115 ms after stimulus onset; Materials and methods). Thus, at the individual neuron level, we replicated the previous results (11), but this time using the linear fits of Equation 1.

**Figure 2.**
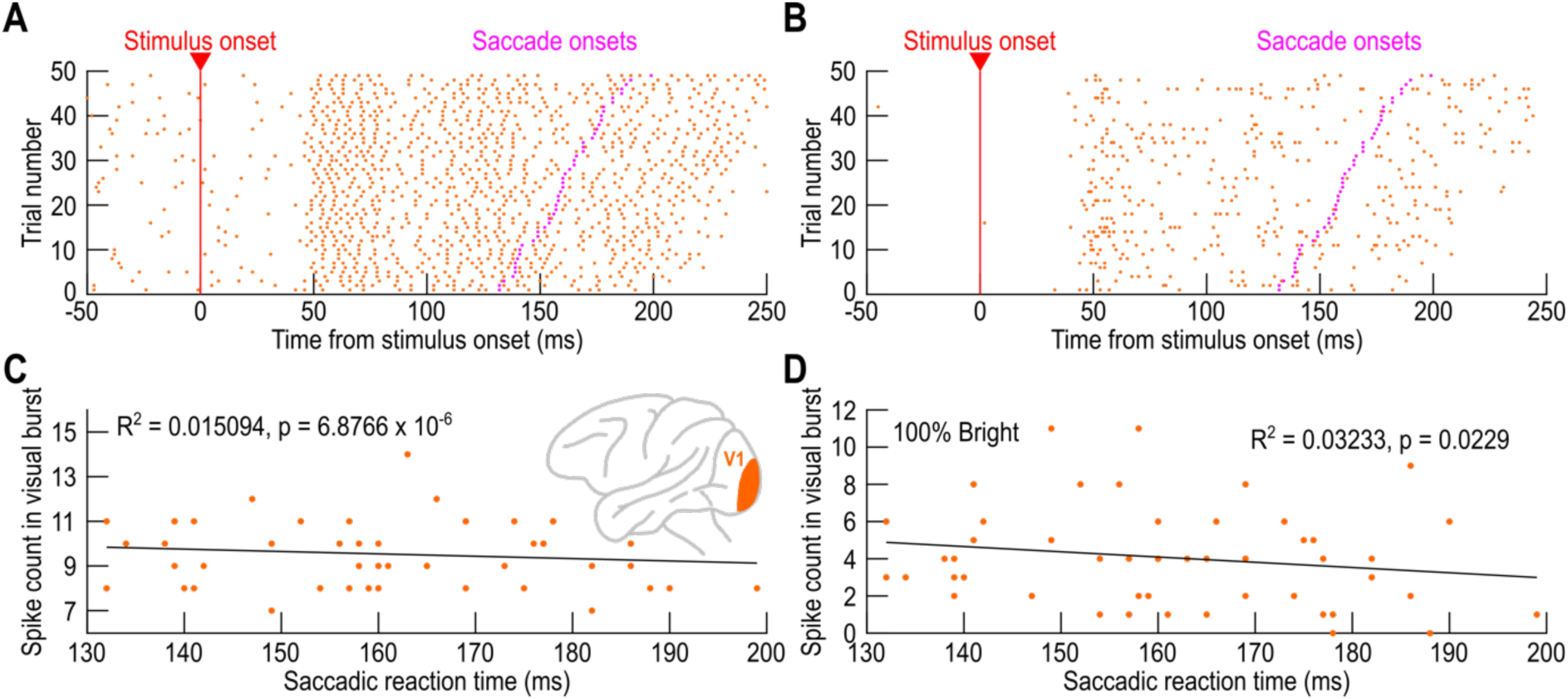
Simultaneously recorded V1 neurons were less related to saccadic reaction time than in the SC. This figure is formatted identically to Fig. 1, except that we now show a pair of simultaneously recorded V1 neurons (from another session), again with 100% contrast bright stimuli as the saccade target. The neurons only emitted a visual response but no saccade-related burst. Importantly, visual response strength did not covary with saccadic reaction time as well as in the case of the pair of SC neurons in Fig. 1.

With more simultaneously recorded neurons within the same SC session, there was a much more dramatic difference between SC and V1 visual responses with respect to predicting saccadic reaction times. Consider, for example, Fig. 3, showing results from the same example SC and V1 sessions of Figs. 1 and 2, respectively. In this figure, we fit the model of Equation 1 (Materials and methods) with successively increasing numbers of simultaneous neurons (*N*) included into the model. For example, for *N* = 2, we picked multiple combinations of pairs of simultaneously recorded neuron within the same session (Materials and methods). We then calculated, for each pair, the goodness of fit of the model (*R^2^_adj_*; Materials and methods). Finally, we plotted the mean and standard deviation of the *R^2^_adj_* value of the session across all tested combinations of simultaneously recorded neuron pairs, and we repeated this for all other *N* values. In the SC (Fig. 3A), there was a monotonic increase in model fit quality as a function of how many neurons were included into the model; the more neurons, the better the prediction of the saccadic reaction time variability became. We confirmed this statistically in two ways. First, a t-test on the ten mean *R^2^_adj_* values against zero revealed a significant result (p<0.0001). Second, we fit the mean values of Fig. 3A with the relationship of Equation 3 (Materials and methods), and we obtained the gray curve in the figure (*a* = 0.0956, *b* = 0.1919 in Equation 3; p<0.0001 for each parameter; *R^2^* = 0.99). Remarkably, for V1 (Fig. 3B), there was no change in the predictive power of the model with more included simultaneously recorded neurons. For example, a linear fit having near-zero slope is shown with the gray curve in Fig. 3B (*a* =-0.0016, *b* =-0.0004 in the version of Equation 3 without the logarithmic transformation of *N*; p = 0.3968 for the intercept, and p = 0.1845 for the slope; R^2^ = 0.2086). Thus, even though we observed short-latency visual responses (from the SC and V1) in both of these sessions (Figs. 1A, B, 2A, B), there was a striking difference in their relationship to saccadic reaction times; this difference was even more compelling than the previous observations at the individual neuron level (11).

**Figure 3.**
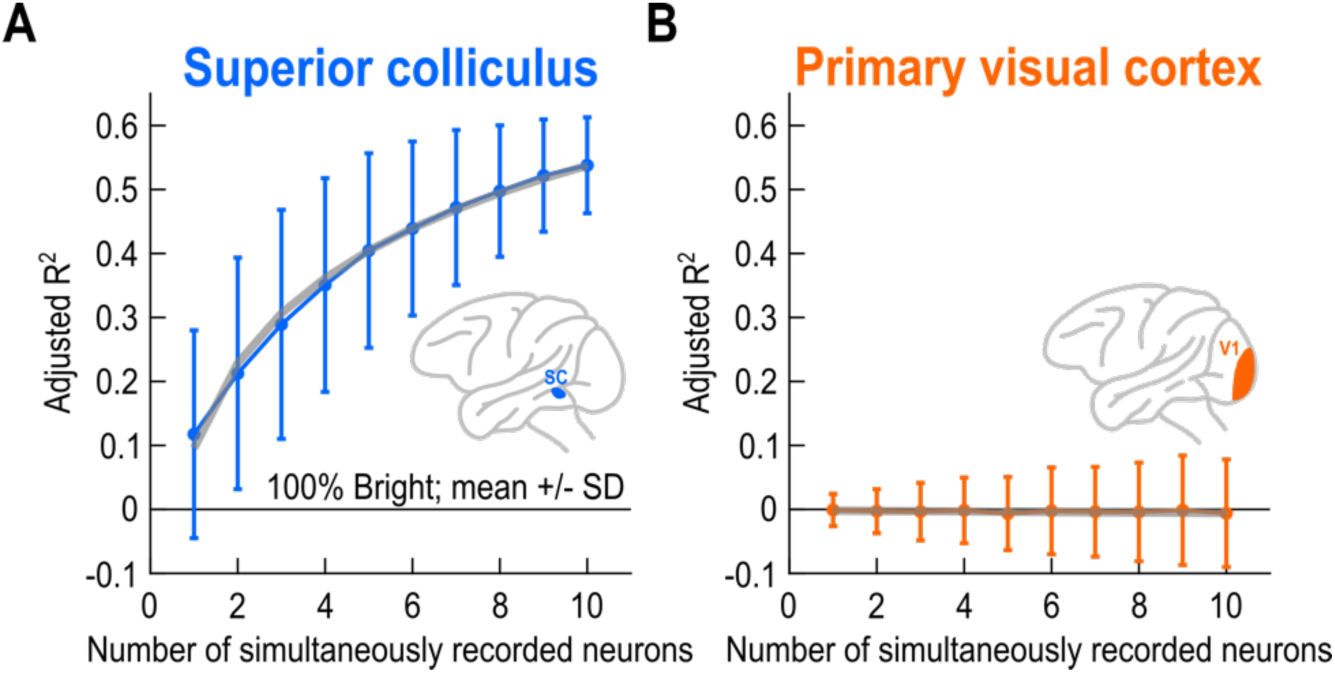
More simultaneously recorded SC neurons improved predictions of saccadic reaction times from visual response strengths, but this was not the case for V1. **(A)** For the same example session and stimuli as in Fig. 1, we plotted the quality of the fit of Equation 1 (*R^2^adj*) as a function of number of simultaneously recorded SC neurons in the model. For each number of simultaneously recorded neurons on the x-axis, we tested the model with different combinations of the neural population (Materials and methods). For example, if we were testing the model with 2 simultaneously recorded neurons (and since we recorded more than 2 neurons in this session), we tested the model with different combinations of pairs of neurons. The error bars denote the standard deviation across different neuron group permutations. Adding more neurons to the model of Equation 1 clearly improved the prediction of saccadic reaction time from SC visual response strength. This is also indicated by the gray curve, which shows the fit of the ten mean values to Equation 3 (Materials and methods); there was a clear monotonic increase. **(B)** For the same example V1 session of Fig. 2, this monotonic increase was not the case. The gray line shows the linear fit of the ten mean values (like Equation 3 but without the logarithmic transformation of *N* in the x-coordinate), indicating no improvement in the prediction of saccadic reaction time with the addition of more simultaneously recorded neurons. Other than the brain area, this panel is formatted identically to **A**.

In our experiments, we also tested different contrasts of the bright luminance polarity. We did so because stimulus contrast can potentially alter the reliability of visual responses (43), leading to the question of whether links to behavior can get modified at different contrast levels. Indeed, in some earlier studies (8), relatively low contrasts were used. Across all sessions, and across all different contrasts, the above results (like Fig. 3) were consistently observed. To summarize them, we obtained the average *R^2^_adj_* curve as a function of the number of simultaneously recorded neurons within a given session (for example, the mean values for the individual sessions in Fig. 3). Then, we averaged all such curves across all sessions, within either the SC or V1. The results in the SC always showed an increase in model fit quality (*R^2^_adj_* when fitting Equation 1) as a function of neuron number (Fig. 4; each panel shows one stimulus contrast level, and the error bars denote SEM across sessions). In stark contrast, the V1 curves were always around zero, regardless of neuron number.

**Figure 4.**
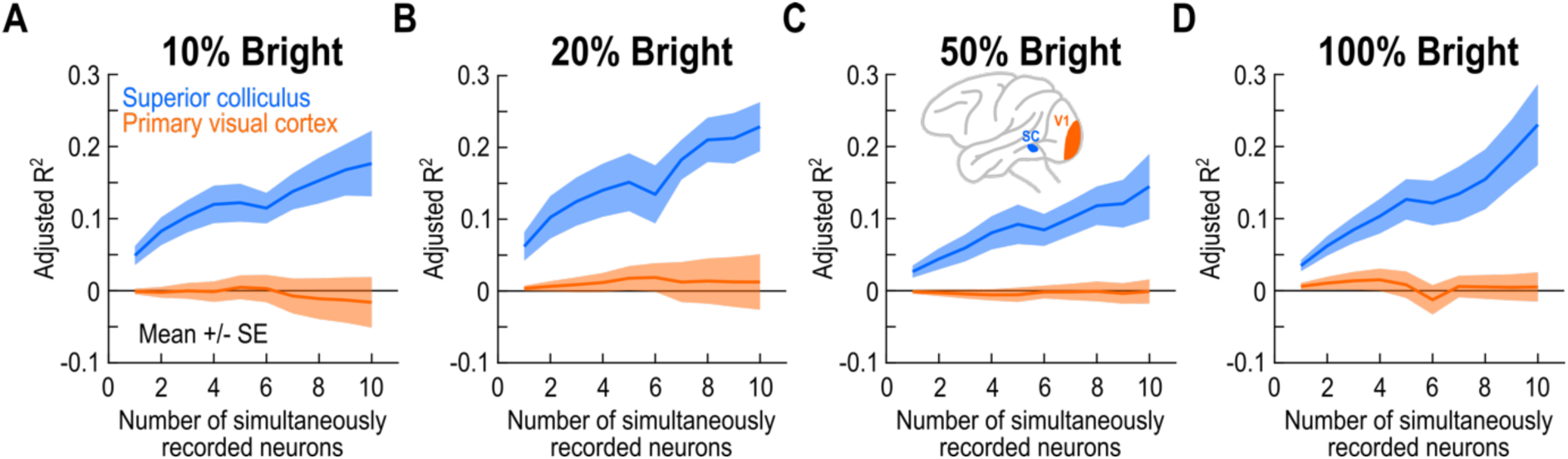
The results of Figs. 1-3 were consistent across sessions in both brain areas. **(A)** Each curve shows the average across sessions (28 for the SC and 13 for V1) of curves like those in Fig. 3, but only for the case of the saccade target being a bright stimulus of 10% contrast. Error bars denote the standard error across sessions. In the SC, model predictions of saccadic reaction time (from visual response strength) consistently improved by increasing the number of simultaneously recorded SC neurons. This was not the case in V1. **(B-D)** Similar results for the other stimulus contrasts of our bright stimuli. In all cases, the V1 predictions always remained at an *R^2^_adj_* value near 0.

We confirmed these observations statistically by running a 3-way ANOVA with brain area, contrast, and number of neurons as the factors. There was a main effect for all factors (F = 345.58, p<0.0001 for brain area; F = 8.47, p<0.0001 for contrast; and F = 8.13, p<0.0001 for number of neurons). Importantly, note how the SC curves (blue in Fig. 4) always started above zero at N=1, which is another way of demonstrating the single neuron results of the previous report (11). Indeed, at each contrast level individually, the SC *R^2^_adj_* values were greater than zero (p<0.0001 for each contrast level; t-test for being larger than zero). On the other hand, in the case of V1, the results for individual contrast levels were not systemtically larger than zero (p = 0.7395 for 10%, p = 0.0344 for 20%, p = 0.8605 for 50%, and p = 0.0701 for 100%; t-test for being larger than zero). Thus, our observations indicate that even when considering multiple concurrently active neurons after stimulus onset, SC visual responses are much better correlated with trial-by-trial saccadic reaction time variability than V1 visual responses.

### Dark contrasts are different in the primary visual cortex

Our experiments also included stimuli of negative luminance polarity (darker than the surrounding gray background; Materials and methods) (11). At the single neuron level, our previous report did not find a substantial qualitative difference between these stimuli and the bright contrasts in terms of the links between their associated neural visual responses and saccadic reaction times (11). However, using simultaneous neuron analyses, here we found that 100% contrast dark stimuli were different from other stimuli in the case of V1. To demonstrate this, we repeated the above analyses but for the dark stimuli.

In the SC, we obtained qualitatively similar results regardless of luminance polarity. For example, Fig. 5A-C shows results for the 100% contrast bright stimuli from an additional example session in our database (formatted like in the first example session of Figs. 1, 3A), and Fig. 5D-F shows the results from the same session with 100% contrast dark stimuli. In Fig. 5A, D, the visual responses of one neuron to either the bright (Fig. 5A) or dark (Fig. 5D) stimuli exhibited a negative correlation with saccadic reaction time. This was also the case in Fig. 5B, E for another simultaneously recorded neuron from the same session (again for the two types of stimuli). Across populations of simultaneously recorded neurons within the same session, a similar monotonic increase in *R^2^_adj_* value was evident for both luminance polarities (Fig. 5C, F). Statistically, a t-test on the ten mean *R^2^_adj_* values in Fig. 5C revealed a significant difference from zero (p<0.0001; tested for being larger than zero), and this was also the case in Fig. 5F (p<0.0001; t-test for being larger than zero). Similarly, a regression along Equation 3 revealed clear monotonic increases (*a* = 0.0584, *b* = 0.1183 for Fig. 5C; p<0.0001 for both parameters; R^2^=0.99; and *a* = 0.0439, *b* = 0.0783 for Fig. 5F; p<0.0001 for both parameters; R^2^ = 0.99). Thus, the results of Figs. 1-4 were replicated for dark contrasts in the SC, at least in this example session.

**Figure 5.**
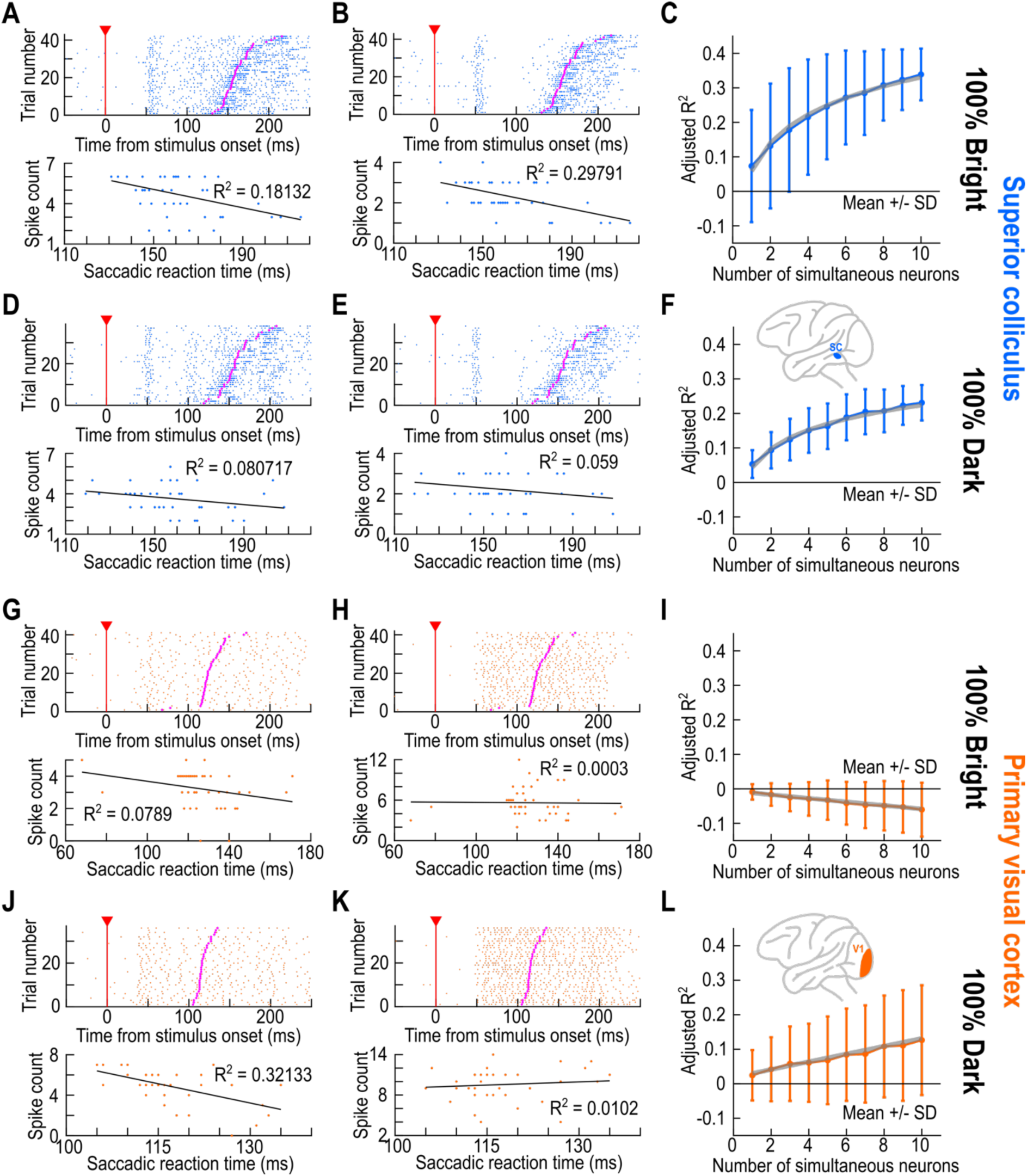
High contrast dark stimuli were different in V1. **(A)** The responses of an example SC neuron formatted in a similar fashion to Fig. 1A, C. The neuron exhibited a negative correlation between visual response strength and saccadic reaction time. **(B)** Similar observations from a second SC neuron recorded from the same session. **(C)** Across all neuron combinations up to *N* = 10 in Equation 1, the results of this session were similar to those of Figs. 1, 3 for 100% contrast bright stimuli. That is, there was an increase in the quality of predicting saccadic reaction times from the visual response strengths of the simultaneously recorded SC neurons in this session. Error bars denote standard deviation across neuron permutations for each *N* (Materials and methods), and the gray line is the fit to Equation 3. **(D)** The same neuron as in **A** showed similar dependencies between visual responses and saccadic reaction times for a 100% contrast dark stimulus as the saccade target, but the correlation strength (bottom panel) was slightly weaker. **(E)** Similar observations for the same neuron of **B**, recorded from the same session. **(F)** Across neuron combinations of the same session, there was again a monotonic increase in prediction quality for the dark stimuli with increasing numbers of simultaneously recorded neurons. **(G-I)** Results from an example V1 session and 100% contrast bright stimuli. This session replicated the observations of Figs. 2, 3. Note that the gray line in **I** is the fit to Equation 3, but without the logarithmic transformation of the number of neurons (Materials and methods). Also note that there was no monotonic increase in **I**, but a decrease instead. **(J-L)** For 100% contrast dark stimuli, however, the V1 session exhibited increased prediction quality with more neurons included in the model of Equation 1. Specifically, there was a monotonic linear increase with increasing numbers of simultaneously recorded neurons, which was absent for the bright stimuli.

For V1, the example session of Fig. 5G-L revealed that adding more neurons to the model of Equation 1 actually improved the prediction quality of saccadic reaction times when the stimulus was a 100% contrast dark disc (Fig. 5L). This effect was different from that for the 100% contrast bright stimulus in the same session (Fig. 5I), in which the model was incapable of capturing similar trends in the data even with 10 simultaneously recorded neurons. In fact, the negative slope of Fig. 5I indicates that our model was even worse than a lack of linear relationship between visual responses and saccade times. Moreover, the individual example neurons in this session (Fig. 5G, H, J, K) showed generally weaker *R^2^_adj_* values than for the SC neurons. We again confirmed these observations statistically. For the bright contrast (Fig. 5I), the V1 neuron showed a negative slope in the linear fit of Equation 3 (without the logarithmic transformation of *N*; Materials and methods), but the neuron showed a positive slope for the dark contrast (for the bright contrast: *a* =-0.0066, *b* = - 0.0052; p = 0.0020 for *a*; p<0.0001 for *b*; R^2^ = 0.9834; for the dark contrast: *a* = 0.0187, *b* = 0.0113; p = 0.0005 for *a*, p<0.0001 for *b*; R^2^ = 0.9825). Consistent with this, a t-test of the 10 *R^2^_adj_* values showed significance for the measurements being smaller than zero in the bright contrast (p<0.0001), and a significance for the measurements being larger than zero in the dark contrast (p<0.0001).

Across sessions with the dark luminance polarity, there was a clear increasing relationship in *R^2^_adj_* value in V1 with increasing numbers of simultaneously recorded neurons at 100% contrast (Fig. 6); this effect was missing for bright contrasts (Fig. 4; the data of Fig. 4 are shown in faint colors in Fig. 6 for easier direct comparison). Statistically, at each contrast level individually, both the SC and V1 *R^2^_adj_* values in Fig. 6 were greater than zero (p<0.0001 for each contrast level; t-test for being larger than zero). Similarly, a 3-way ANOVA on the data in Fig. 6 revealed significant main effects of brain area (F = 264.46, p<0.0001), contrast (F = 12.12, p<0.0001), and number of simultaneously recorded neurons (F = 14.25, p<0.0001). Post-hoc inspection of the figure revealed the highest positive slope in the orange curves for the 100% contrast condition.

**Figure 6.**
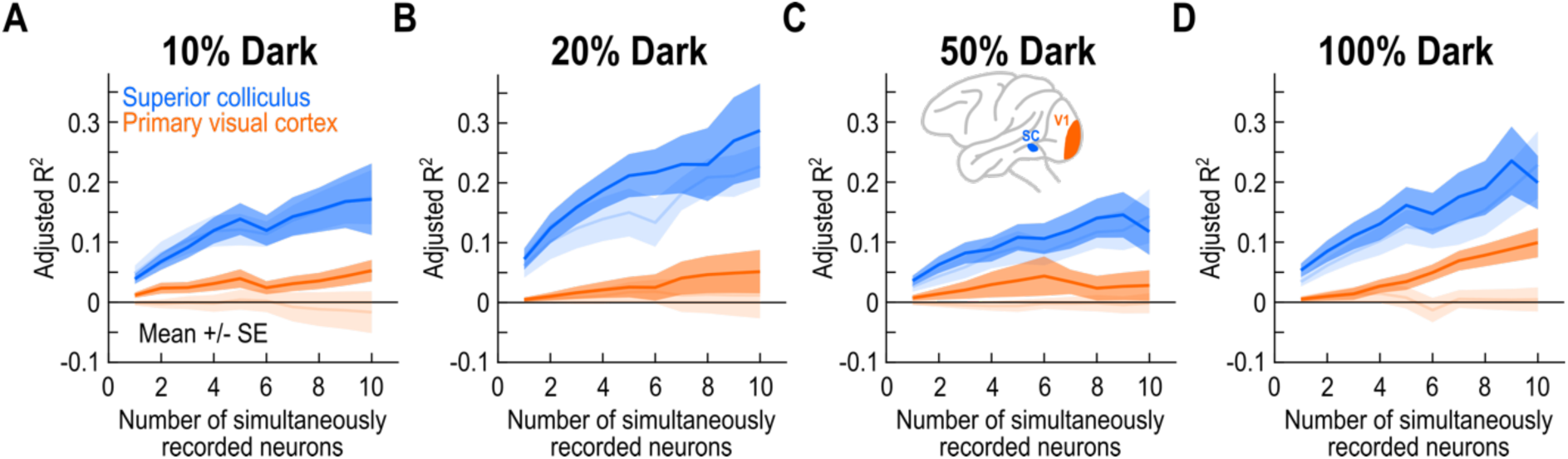
Similar to Fig. 4 but with dark contrasts; 100% contrast dark stimuli behaved differently for V1 neurons. This figure is formatted identically to Fig. 4. At 100% contrast, there was a clear improvement in saccadic reaction time predictability based on V1 visual response strengths (orange curve). Note that at the individual neuron level (1 on the x-axis of **D**) V1 was not a good predictor at all with 100% dark contrasts. Thus, this improvement with more neurons was a distinct property of the V1 population, which could not be predicted based on the results of the previous study alone (11). The faint curves show the same data of Fig. 4 for easier comparison.

It is also interesting to note here that at the single neuron level, there was no appreciable difference in model fits between 100% contrast bright or dark stimuli (see the *N* = 1 data points in Figs. 4D, 6D; the curves for brights and darks are also shown together in Fig. 6D for easier comparison). This was also the case in the previous report (11). For 10% dark stimuli, in particular, the model fit quality remained higher than zero in V1 across neuron counts (Fig. 6A; orange curve), but without as strong a positive slope as for 100% dark stimuli (Fig. 6D; orange curve); this effect at 10% contrast could be attributed to monkey F’s data, as we observed previously for this monkey at the single neuron level (11). However, at 100% contrast, none of the monkeys individually showed effects similar to Fig. 6D at the single neuron level (11). Thus, with simultaneously recorded neurons, the clearest evidence for a link between V1 visual responses and saccadic reaction times emerged for 100% contrast dark stimuli in our database; and, this evidence could not be predicted from the single neuron analyses.

Finally, we performed a multi-factor ANOVA on the aggregate data, and we found significant main effects of contrast (F = 19.69, p < 0.0001), luminance polarity (F = 28.17, p < 0.0001), number of simultaneously recorded neurons (F = 21.98, p < 0.0001), and brain area (F = 607.32, p < 0.0001). Importantly, the factor brain area had the biggest effects, indicating a substantial difference between V1 and SC visual responses in the prediction of saccadic reaction times.

### Stronger effects in superior colliculus visual-motor than visual-only neurons

We next asked whether the weakest possible SC correlations to behavior could be as weak as those that we observed with V1 above. Specifically, prior work has shown that visual responses in SC visual-motor neurons reflect reaction times better than visual responses of purely visual SC neurons (10–12). This prompted us to explore this idea further with our simultaneously recorded neurons. Figure 7 shows SC results in analyses like those shown in Figs. 4, 6 above, but after separating the visual-motor and visual-only SC neurons. Note that because we now had smaller numbers of neurons in each group, we restricted our analyses to a limit of 5 simultaneously recorded neurons of each type (Materials and methods). The figure also shows, in faint colors, the original results of Figs. 4, 6 for easier comparison. As can be seen, the curves for visual-motor neurons were always higher than those for visual-only neurons, and they also generally exhibited sharper increases with increasing numbers of simultaneously recorded neurons. Thus, we confirmed that the visual responses of visual-motor neurons are the best predictors of saccadic reaction times in the SC.

**Figure 7.**
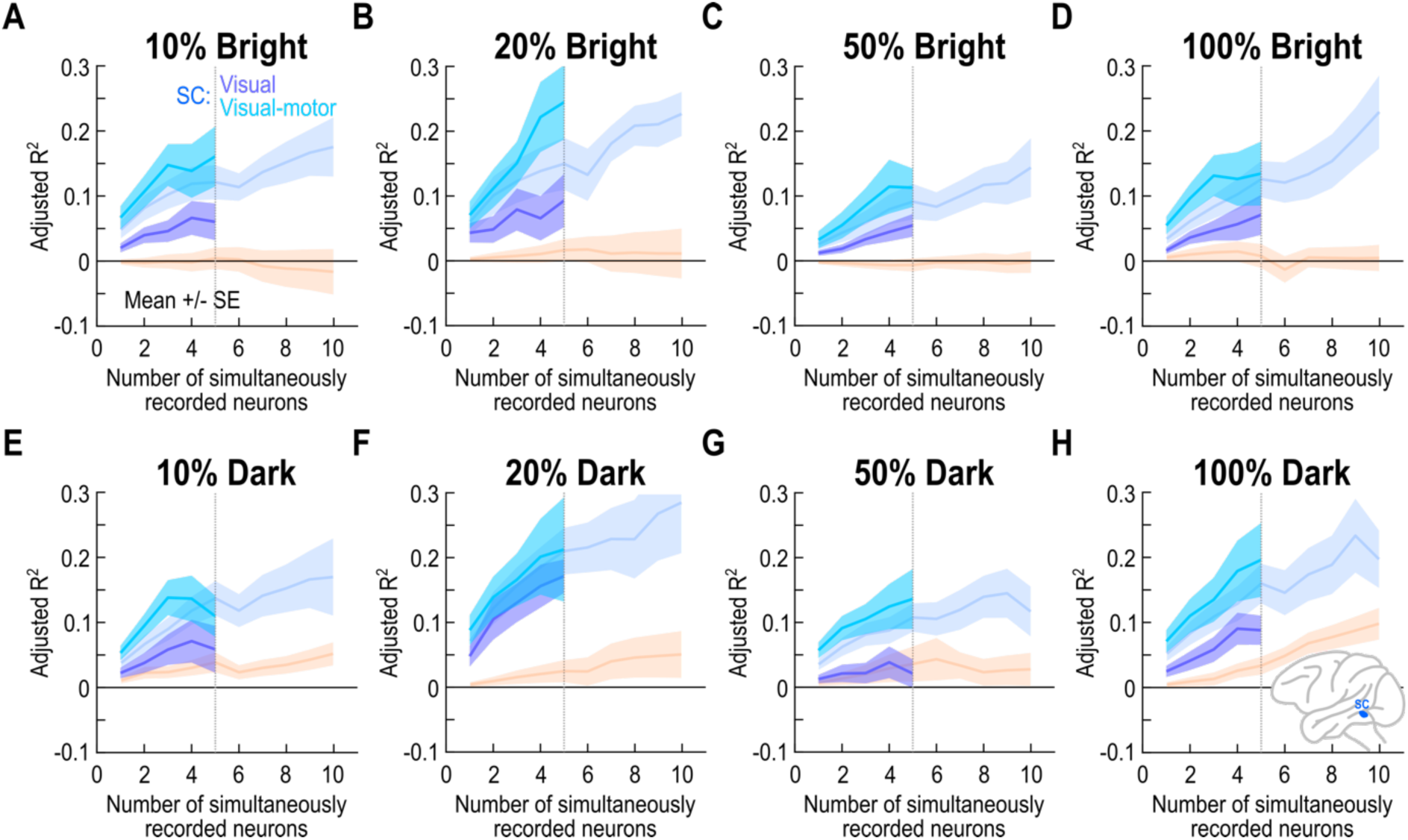
Visual-motor SC neurons showed the best relationship to saccade timing. (A-D) Results like those shown in Fig. 4, but after separating SC neurons into visual-only and visual-motor functional types (Materials and methods). The faint colors show the global SC (faint blue) and V1 (faint orange) results from Fig. 4 for easier comparison. Also note that since we were dealing with smaller numbers of visual-only or visual-motor neurons per session than in the aggregate population, we restricted our analyses to 5 simultaneously recorded neurons. This was still sufficient to show the levels and slopes of the different curves. As expected from prior work (11, 12), visual-motor neurons showed the best relationship to saccade timing. Note also how the slopes of the visual-motor curves were generally higher than those for the visual-only neurons. **(E-H)** Similar results but for the dark contrasts. Note how even the visual-only curves were practically always higher than the V1 curves (for either bright or dark contrasts), suggesting that even visual-only neuron visual responses are better related to saccade timing than V1 visual responses (11). All other conventions are like in Figs. 4, 6. Error bars denote SEM, and the same number of sessions was included in this figure as in Figs. 4, 6.

Interestingly, SC visual-only neurons were still better than V1 neurons in their relationship to saccadic timing, despite having weaker effects than visual-motor neurons. In fact, even for 100% dark stimuli (Fig. 7H), the curve for SC visual-only neurons was higher than the curve for V1 neurons. Therefore, even though the visual-only neurons are the biggest recipients of early cortical visual input (19, 38–42), these neurons do not copy their V1 inputs in terms of the relationship to active orienting behavior.

Statistically, we ran a multi-factor ANOVA with contrast, luminance polarity, number of simultaneously recorded neurons, and SC functional neuron type (visual-only or visual-motor) as the factors. There were significant effects for all factors except for luminance polarity, consistent with Fig. 7 (F=17.05, p<1×10^-30^ for contrast; F=24.51, p<1×10^-30^ for number of neurons; F=137.56, p<1×10^-30^ for SC functional cell type; F=3.56, p=0.0595 for luminance polarity).

Therefore, even within the SC itself, visual responses of different functional SC neuron types can be differentially related to the timing of saccadic eye movements; but, the effects are always still stronger than the V1 effects.

## Discussion

In this study, we asked to what extent could V1 visual responses be linked to trial-by-trial saccadic reaction time variability when multiple concurrently activated neurons are taken into account. Our motivation was that we recently found much higher covariation in SC than V1 visual responses with such eye movement timing variability (11). However, this was only done at the individual neuron level. Here, we analyzed multiple simultaneously recorded SC or V1 neurons within a given session. We linked the visual response strengths of such simultaneous neurons to saccadic reaction time with as simple a model as possible. We found that the predictive power of SC visual response strengths, in terms of saccade timing, increased monotonically with the addition of more simultaneously recorded neurons. Remarkably, and especially for positive luminance polarity stimuli, this was not the case in V1, with the predictive power remaining at zero even with up to ten simultaneously recorded neurons (Fig. 4). Only with 100% contrast dark stimuli was there the clearest benefit to adding more V1 neurons in predicting saccadic reaction times (Fig. 6). We believe that these observations highlight the fundamental functional difference that exists between SC and V1 visual responses, despite their qualitative similarity (19).

Our SC results add to convincing evidence that SC visual responses are intricately related to saccade timing (9–19). These results also show that such intricate relationship is strongest for visual-motor neurons (Fig. 7) consistent with our previous observations (11). The primary reason that we included all SC neurons in the some of our other analyses (such as those in Figs. 4, 6) is that we wanted to maximize the size of our neural database. However, the separation of SC neurons into two functional types also allowed us to see whether V1 predictions could approach even the lower bound of SC predictions (namely, those of the visual-only neurons). This was clearly not the case (Fig. 7), cementing the interpretation that SC visual responses exhibit much higher covariation with saccadic reaction time variability than V1 visual responses (11).

It would be interesting in the future to investigate the implications of our observations on the links between pre-stimulus cortical state and saccadic timing (44, 45). For example, saccades synchronize and interact with occipital alpha rhythms (46, 47), and subsequent saccade timing can depend on the previously phase-reset oscillatory rhythms in behavior (48). This makes it important in subsequent studies to investigate pre-stimulus state in both the SC and V1, and to relate such state to visually-guided saccadic reaction times.

One additional remarkable observation from our present analyses was that dark stimuli, particularly with 100% Weber contrast, were a clear exception in terms of our V1 results. For all bright contrasts, V1 neurons showed essentially no difference from zero in terms of their *R^2^_adj_* values for the model fits (Fig. 4). This is a noteworthy difference from previous predictions from V1 neurons (8), although the stimuli and tasks were different in this earlier study. On the other hand, with dark stimuli (Fig. 6), V1 neurons did show some predictive power of saccade timing. For 10% contrasts (Fig. 6A), this was likely driven by monkey F, since our previous individual monkey analyses revealed that this monkey (with exceptionally fast reaction times) had clearer predictive power of V1 visual response strength for saccade timing at 10% contrasts (for dark stimuli) (11). Surprisingly, at 100% contrast levels, no predictions from the previous study could explain the present study’s observations (Fig. 6D): for 100% contrast dark stimuli, there was indeed a benefit to adding more V1 neurons to the model predictions; the results with *N* = 1 were similar to our previous study. This population phenomenon adds to the value of our present study’s contributions, because it represents a result that could not have been easily predicted from only the single neuron analyses. This population phenomenon could also reflect the preference of V1 neurons for dark stimuli. Indeed, most V1 neurons do prefer dark over bright contrasts (49), and there are other cortical asymmetries related to dark stimuli as well (50–55). While some asymmetries between dark and bright stimulus processing do exist in the SC as well, particularly for visual response timing (17), the numbers of SC neurons preferring bright or dark stimuli seems to be a bit more balanced than in V1, although a direct comparison within the same animals and with the same stimuli and displays remains to be performed. Thus, it could be that the ecological relevance of dark stimuli in natural scenes (49–55) favor having a better link to active behavior in V1 for these stimuli, when compared to bright ones.

Of course, it remains possible that a more complicated readout of V1 neurons could still be used to predict saccadic reaction time from V1 visual responses. However, the stark difference that we observed between V1 and the SC (with our simplistic linear readout mechanism) implies a much more direct link between SC visual responses and the behavioral output. This makes sense anatomically because the SC’s downstream brainstem projections are causal to driving orienting behaviors (10). Thus, variance in SC visual responses is expected to also have a greater impact on such behaviors than variance in V1 visual responses.

More broadly, our results fit into a wider framework for what dictates saccade timing. On the cognitive and motor sides, there has been significant progress made, with evidence again implicating the SC in such processes. For example, target probability can influence SC motor preparation states (20–22), and thus subsequent saccade generation times. Similarly, the classic SC motor burst itself may be considered a trigger for saccades; as a result, correlations between SC motor burst times and saccade onset times are much higher (23, 24) than correlations of earlier sensory and cognitive processes, including our own observations on visual responses. Having said that, the correlations of visual responses to saccade timing are not trivial at all: before any of the cognitive and motor factors that we just mentioned could come into play, sensing stimuli needs to be completed, and this is where our work fits within this global framework. In fact, in natural viewing scenarios, visual reafferent responses would be expected after every saccade, and these reafferent responses jumpstart the subsequent cognitive and motor processes. Thus, variability in fixation durations in natural scene viewing should depend on sensory factors, and there is indeed evidence supporting that (14).

This leads to the question of whether our stimuli were somehow not optimal for V1 neurons. This is not the case, because a majority of V1 neurons actually do signal luminance polarity (49). Also, in our recent work, we found very robust visual responses in V1 to exactly our types of stimuli (11). More importantly, using Gabor gratings in the SC, we still found remarkable correlations between SC visual responses and saccade timing (14). Thus, even with stimuli that are optimal for V1, such as Gabor gratings, the links between SC visual responses and saccade times were very obvious. Even more recently, we also repeated the experiments of (14) in both the SC and V1 (again, on the same animals and with the same stimuli), and we found a large difference between the two brain areas: the timing dependencies of SC visual bursts as a function of spatial frequency, which were directly reflected in saccadic reaction times and fixation durations in our previous study (14), were categorically not the same in V1 (56). Thus, even with V1 optimal stimuli, like Gabor gratings, the stronger links of the SC to behavior persist. This indicates that anatomical connectivity alone (for example, from V1 to the SC’s visual-only neurons) does not necessarily explain function, and that is why physiological investigation is necessary. A similar sentiment, this time related to retino-tectal projections (41), can be made when considering visually-driven ocular reflexes (57).

Ultimately, it would also be interesting to contemplate visual responses in the context of delayed saccades, whether visually or memory-driven. In such saccades, the instruction to trigger the saccade is temporally dissociated from the time of the visual bursts. In recent related work, We found that in such a situation, a relevant component of the drive for the saccade is a transient signal that appears in foveal SC neurons, and that is triggered by the go instruction for generating the saccade (58). However, we still do not know if there is an interaction between this foveal transient signal and the peripheral visual bursts when it comes to saccadic reaction times. So, this would be interesting to test. Interestingly, we also have behavioral evidence from humans and monkeys, showing that saccadic reaction times in memory-guided saccades can reflect saccade direction (6) – specifically, being shorter for upward saccades. Given that SC visual responses are stronger for upper visual field locations (18), this is suggestive that there could indeed be a relationship between visual burst strength and saccadic reaction time even in memory-guided saccades as well.

## Grants

We were funded by the Deutsche Forschungsgemeinschaft (DFG; German Research Foundation) through the following projects: SPP 2411 Sensing LOOPS: Cortico-subcortical Interactions for Adaptive Sensing (project numbers: 520617944 and 520283985, HA 6749/11-1); SFB 1233 Robust Vision (project number: 276693517); BO5681/1-1.

## Author contributions

Conceived and designed research: YY, ZMH

Analyzed data: YY, ZMH

Interpreted results of experiments: YY, ZMH

Prepared figures: YY, ZMH

Drafted manuscript: YY, ZMH

Edited and revised manuscript: YY, ZMH

Approved final version: YY, ZMH

